# Origin and development of uniparental and polyploid blastomeres

**DOI:** 10.1101/2024.07.30.605883

**Authors:** Yan Zhao, Andrea Fernández-Montoro, Greet Peeters, Tatjana Jatsenko, Tine De Coster, Daniel Angel-Velez, Thomas Lefevre, Thierry Voet, Olga Tšuiko, Ants Kurg, Katrien Smits, Ann Van Soom, Joris Robert Vermeesch

## Abstract

Whole-genome (WG) abnormalities, such as uniparental diploidy and triploidy, cause fetal death. Occasionally, they coexist with biparental diploid cells in live births. Understanding the origin and early development of WG abnormal blastomeres is crucial for explaining the formation of androgenotes, gynogenotes, triploidy, chimerism, and mixoploidy. By haplotyping 118 blastomeres from first cleavages, we identified various mechanisms of heterogoneic divisions that lead to WG abnormal blastomeres or their coexistence with normal blastomeres in both multipolar and bipolar cleaving zygotes. After culturing the totipotent blastomeres to three preimplantation stages and performing transcriptome profiling on over 600 cells, we discovered that stress responses contribute to developmental impairment in WG abnormal cells, resulting in either cell arrest or blastocyst formation. Some first-cleavage-derived WG abnormal blastomeres can survive early development and progress to blastocysts. Their potential dominance in preimplantation embryos represents an overlooked cause of abnormal development. Haplotype based screening could further increase pregnancy rates.

## Introduction

Chromosomal abnormalities are common during human development, especially in the preimplantation phase(Hassold et al., 1980; Hassold and Hunt, 2001; Vanneste et al., 2009). One notable category of chromosome abnormalities are whole-genome (WG) anomalies(Masset et al., 2021). Embryos bearing WG abnormalities contain abnormal levels of parental genomes rather than the normal diploid constellation with one maternal and one paternal haploid genome. For instance, uniparental haploid or diploid embryos carry only chromosomes from a single parent, while triploid embryos contain an additional haploid set of chromosomes from one of the parents. Constitutional WG anomalies are generally lethal during the embryonic stage. Triploidy occurs in approximately 1% of all conceptuses, contributing to around 10% of all spontaneous abortions(Hassold et al., 1980; Zaragoza et al., 2000) with extremely rare live born cases surviving less than a year(Sherard et al., 1986). Complete and partial hydatidiform moles, occurring at rates of 1 in 1000 and 3 in 1000 pregnancies respectively, are primarily androgenetic diploid and diandric triploid, respectively(Seckl et al., 2010). Mosaic forms with coexistence of normal cells can result in rare cases of live birth. For example, gynogenetic and androgenetic chimerism (coexistence of uniparental and normal cells) and diploid/triploid mixoploidy (coexistence of triploid and normal cells) have been reported in live-born individuals with congenital abnormalities(Madan, 2020). As a model organism with a developmental process and an aneuploidy profile similar to that of humans(Hansen, 2010; Santos et al., 2014; Tšuiko et al., 2017), WG abnormalities have also been identified at various stages during bovine development(De Coster et al., 2022; Destouni et al., 2016; Dunn et al., 1970; Meinecke et al., 2003; Tšuiko et al., 2017).

Several models have been proposed to explain the mechanistic origins of WG abnormalities in embryos(Masset et al., 2021). These include dispermy or diploid oocytes leading to triploidy, as well as parthenogenetic division of the oocyte or fertilization of an ‘empty’ egg resulting in uniparental embryos. While these models explain the origin of triploid or uniparental embryos, they are limited by the deduction from surviving cells at later developmental stages. Additionally, the mechanistic origins of chimerism and mixoploidy remain speculative(Madan, 2020; Masset et al., 2021). Since all body cells stem from a single zygote, diverse WG abnormalities observed during the late developmental stages are likely the result of irregular zygotic cleavages. Our recent study with bovine *in vitro* fertilized embryos provides direct evidence of a non-canonical first zygotic division as the mechanistic origin of cell lines exhibiting WG abnormalities(De Coster et al., 2022), which we termed “heterogoneic division”. This specialized division involves the segregation of entire parental genomes into distinct blastomeres, often triggered by polyspermy and occurring concurrently with multipolar zygotic division. Embryos following heterogoneic division typically contain blastomeres with diverse chromosomal constitutions, including biparental diploid, polyploid, and uniparental blastomeres, resembling those identified in chimeric and mixoploid individuals. Such cleavage involving WG segregation errors has also been reported in *in vitro* human and non-human primate embryos(Daughtry et al., 2019; Ottolini et al., 2017).

Heterogoneic division, as the natural origin of blastomeres with WG abnormalities, represents a fundamental yet underexplored aspect of embryo development. While research on preimplantation embryo development often focuses on aneuploidies and their implications(Fernandez Gallardo et al., 2023; Fragouli et al., 2013; Tšuiko et al., 2021), there has been a lack of investigation into the developmental trajectory of spontaneously arising blastomeres carrying WG abnormalities. Examining the subsequent development of blastomeres following both heterogoneic and normal first zygotic divisions will provide valuable insights into the developmental disparities between blastomeres with WG abnormalities and normal blastomeres. Such insights are essential for advancing our understanding of the mechanistic origins of triploidy, moles, chimerism, and mixoploidy identified late during development, and will guide genetic testing and embryo selection strategies regarding WG abnormalities.

Here, we explored the genome constitution and developmental potential of individual blastomeres resulting from both bi- and multipolar first zygotic divisions. Through haplotyping of 118 blastomere outgrowths collected at three preimplantation stages, we demonstrated that uniparental and polyploid blastomeres arise from both multipolar and bipolar cleaving zygotes and exhibit a reduced blastocyst rate. Along with single-cell transcriptome analysis of 446 transcriptomes from 124 blastomere outgrowths, we observed impaired transcriptomic development in blastomere outgrowths carrying WG abnormalities, starting from major embryonic genome activation (EGA). This impaired development is attributed to stress responses induced in blastomere outgrowths with WG abnormalities during EGA. Despite these challenges, some blastomere outgrowths with WG abnormalities successfully managed this stress period and progressed to the blastocyst stage, underscoring their potential significant role in abnormal embryo development.

## Results

### Whole-genome segregation errors occur in multipolar and bipolar first cleavages

To investigate the developmental program of gynogenetic/androgenetic and polyploid blastomeres following spontaneous heterogoneic division, we exploited the totipotency of blastomeres from early cleavage-stage embryos(Johnson et al., 1995; Willadsen, 1980). Daughter blastomeres from 17 bipolar and 35 multipolar cleaved embryos were dissected and individually cultured to either day 2 (T1, 72 hours post fertilization (hpf), 4-6 cell stage), day 4 (T2, 121 hpf, 4-12 cell stage), or day 6 (T3, 170 hpf, blastocyst stage) post blastomere splitting, respectively (**Fig. 1**).

**Figure 1.**
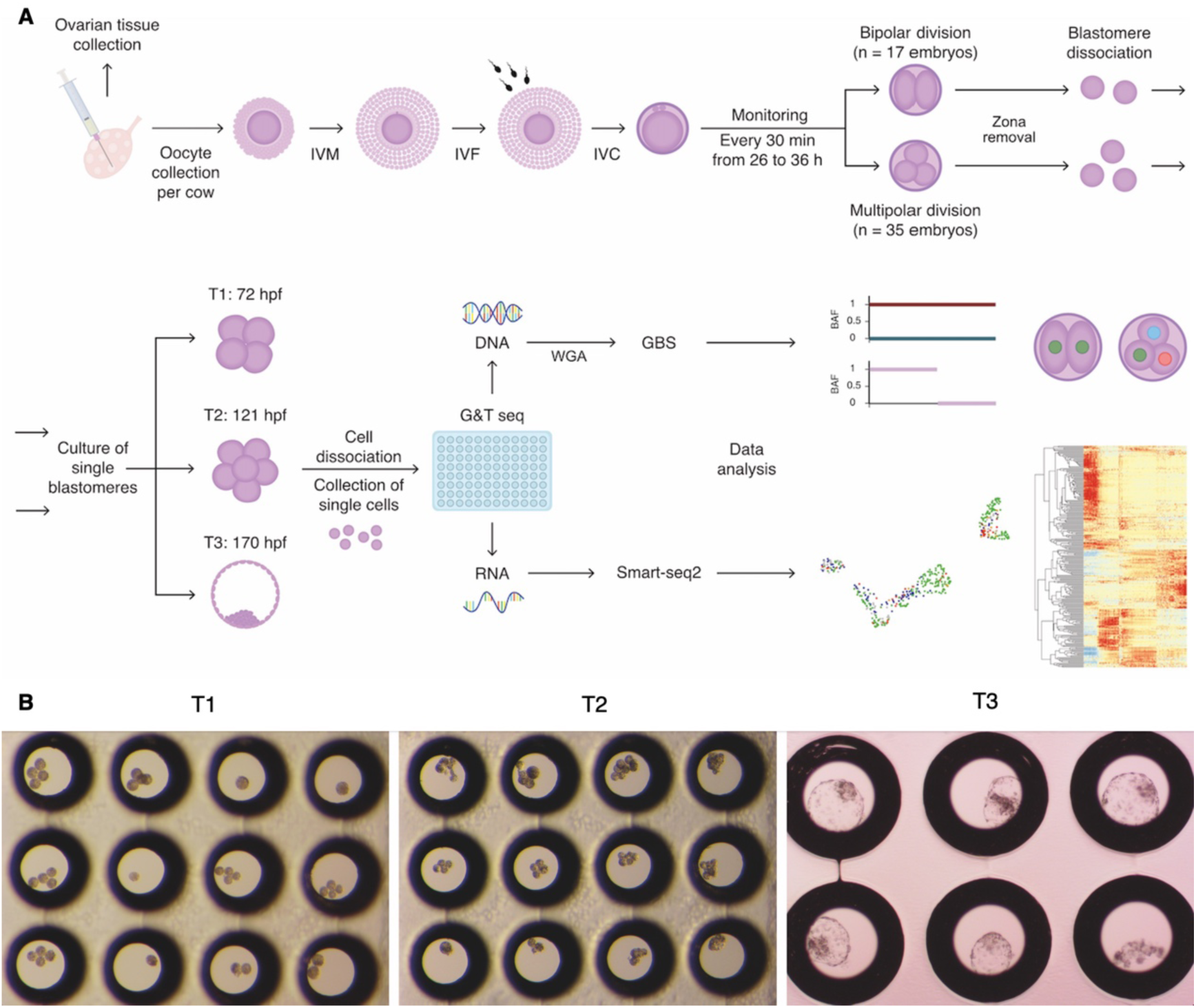
Experimental setup. (A) Diagram illustrating the experimental design. Bovine embryos were created and cultured *in vitro*. After the first zygotic division, blastomeres were dissected and cultured to three preimplantation timepoints. Blastomere outgrowths were subsequently collected and dissected into single cells. Each dissected cell underwent genome and transcriptome separation followed by sequencing. Genome data were used for haplotyping and copy number profiling, while transcriptome data were used for gene expression analysis. Genome and transcriptome sequencing (G&T-seq): Protocol for isolating poly(A) RNA and genomic DNA from single cells. Genotyping-by-sequencing (GBS): Sequencing-based protocol for genome-wide haplotyping and copy number profiling. (B) Example images of blastomere outgrowths collected at T1, T2, and T3, respectively.

To determine the chromosomal constitution of the blastomeres in culture, we performed genome-wide haplotyping and copy number profiling on cells derived from blastomere outgrowths using genotyping-by-sequencing (GBS) (Masset et al., 2022) (**Fig. 1**). We successfully inferred the chromosomal constitution for all cultured blastomeres in 31 embryos and a subset of cultured blastomeres in 12 embryos, totally getting the chromosomal constitution of 118 blastomeres in culture (**Fig. 2A,B; Supplemental Tables S1,S2**). Among these, 45% (n=53) were identified as biparental diploid, while 41% (n=48) exhibited WG abnormalities, including 35 androgenetic, 9 gynogenetic, 3 diandric triploid, and 1 triandric tetraploid blastomeres. Additionally, 14% (n=17) of the blastomeres in culture with inferred chromosomal constitution were found to be anuclear. All anuclear blastomeres and the majority (96%, 46 out of 48) of blastomeres with WG abnormalities originated from embryos undergoing multipolar first cleavage (**Fig. 2B; Supplemental Table S1**).

**Figure 2.**
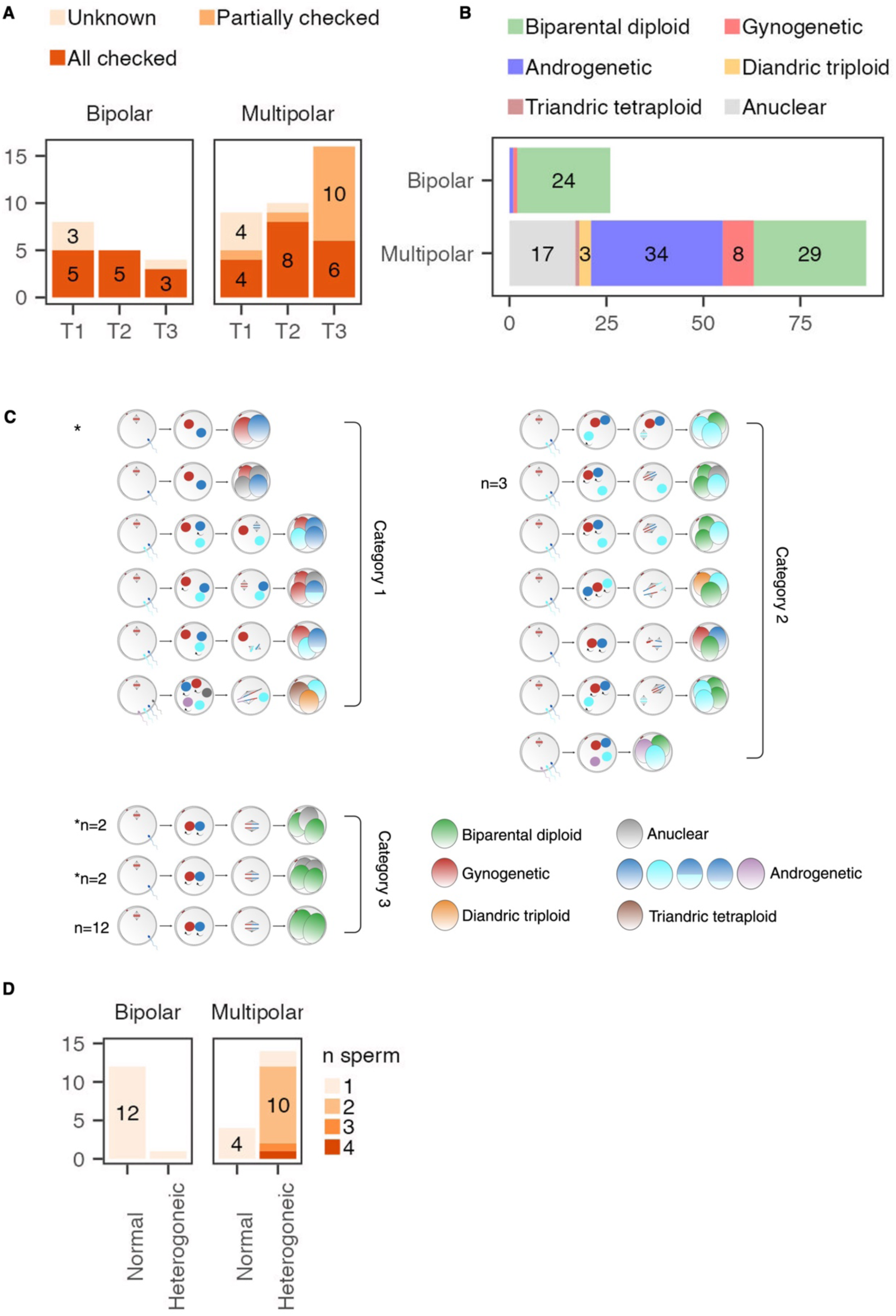
Whole-genome segregation errors occur in multipolar and bipolar first cleavages. (A) Numbers of bipolar and multipolar cleaving embryos used for each blastomere culturing timepoint, categorized based on the completeness of embryo-derived blastomere outgrowth haplotyping: fully haplotyped (All checked), partially haplotyped (Partial), or not haplotyped (Unknown). (B) The inferred chromosomal constitution of 118 blastomeres in culture. (C) Three categories of genome segregation patterns identified during the first cleavage. In the oocyte being fertilized, the second meiotic division is depicted, along with the likely number of sperms fertilizing the embryo as deduced from the haplotypes. The arrows surrounding the pronuclei indicate replication occurred. During the first cleavage, spindles segregating the sister chromatids are shown. In the absence of a spindle, the genome is extruded. Numbers are given for patterns that occurred more than once. * in category 1 indicates the one embryo with bipolar first cleavage and WG segregation error. * in category 3 indicates the four embryos with multipolar first cleavage and without WG segregation error. For androgenetic blastomeres, different colors indicate genetic material from different sperm. (D) Chromosomal segregation patterns for embryos with bipolar or multipolar first cleavage and the inferred number of sperm that fertilized the egg. Normal: Zygote cleaved without WG segregation error. Heterogoneic: Zygote cleaved with WG segregation error. Bipolar and multipolar refer to embryos undergoing either bipolar or multipolar first cleavage, respectively.

For the 31 fully checked embryos, genome composition of all cultured blastomeres allowed us to reconstruct genome segregation patterns during the first cleavage. We observed three broad categories of genome segregation patterns during the first cleavage (**Fig. 2C; Supplemental Table S1**). Category 1 contains 6 embryos with all blastomeres harboring WG abnormalities. Category 2 comprises 9 embryos with a mixture of blastomeres, some with WG abnormalities and others being biparental diploid. Category 3 consists of 16 embryos with only biparental diploid blastomeres. The four multipolar cleaving zygotes in this category also produced anuclear cells. Among the 31 fully examined embryos, 78% (14 out of 18) of those with multipolar first cleavage demonstrated heterogoneic division, while only 8% (1 out of 13) of embryos with bipolar first cleavage showed WG segregation errors (*p*=0.00048) (**Fig. 2C**), confirming the enrichment of heterogoneic division in multipolar cleaving zygotes(De Coster et al., 2022). Furthermore, the high frequency of polyspermic conceptions in embryos undergoing multipolar first zygotic division and exhibiting WG segregation errors(De Coster et al., 2022) was validated, with 86% (12 out of 14) of these embryos inferred to be fertilized by more than one sperm. In contrast, all other embryos with bipolar first cleavage or with multipolar first cleavage but without WG segregation errors were inferred to be fertilized by a single sperm (**Fig. 2D**). Only 10% (1 out of 10) of dispermic embryos give rise to triploid blastomeres, which is consistent with previous observations that dispermic fertilization seldom leads to triploid development(De Coster et al., 2022; Kola et al., 1987). Most of the observed mechanisms driving heterogoneic division align with those from our previous study(De Coster et al., 2022; Destouni et al., 2016). Among the 15 embryos undergoing heterogoneic division, 10 were inferred to involve the extrusion of paternal pronuclei into separate cells, 4 likely involved an independent spindle separating paternal or maternal chromosomes, 2 likely involved a tripolar spindle, and 1 likely involved double spindles (**Fig. 2C**).

Interestingly, two novel genome segregation mechanisms were identified. Firstly, within category 1, we observed an embryo undergoing bipolar first cleavage, yet exhibiting heterogoneic division (**Fig. 2C**). The most plausible explanation is that the two parental zygotic spindles failed to align and instead segregated into distinct blastomeres. Hence, genome-wide segregation errors are not confined to multipolar cleavers. Additionally, within category 3, we identified four embryos undergoing multipolar first cleavage, yet exhibiting normal genome segregation. Specifically, two biparental diploid blastomeres were generated alongside one or two anuclear cells (**Fig. 2C**). These findings suggest that bipolar first cleavage does not always ensure normal genome segregation, and multipolar first cleavage does not inevitably result in WG or chromosomal segregation errors.

### Blastomeres with whole-genome abnormalities can reach blastocyst stage despite impaired developmental potential

To investigate the developmental potential of heterogoneic division-derived blastomeres with WG abnormalities, we compared the developmental stage of the 53 biparental diploid blastomere outgrowths and 48 blastomere outgrowths with WG abnormalities at the time of outgrowth collection (**Fig. 3A,B)**. We observed impaired development for blastomeres with WG abnormalities compared to biparental diploid blastomeres at T3, with 80% (12 out of 15) of biparental diploid blastomere outgrowths developing to blastocysts, while only 21% (6 out of 28) of blastomere outgrowths with WG abnormalities reached this stage (*p*=0.0016) (**Fig. 3B,C**).

**Figure 3.**
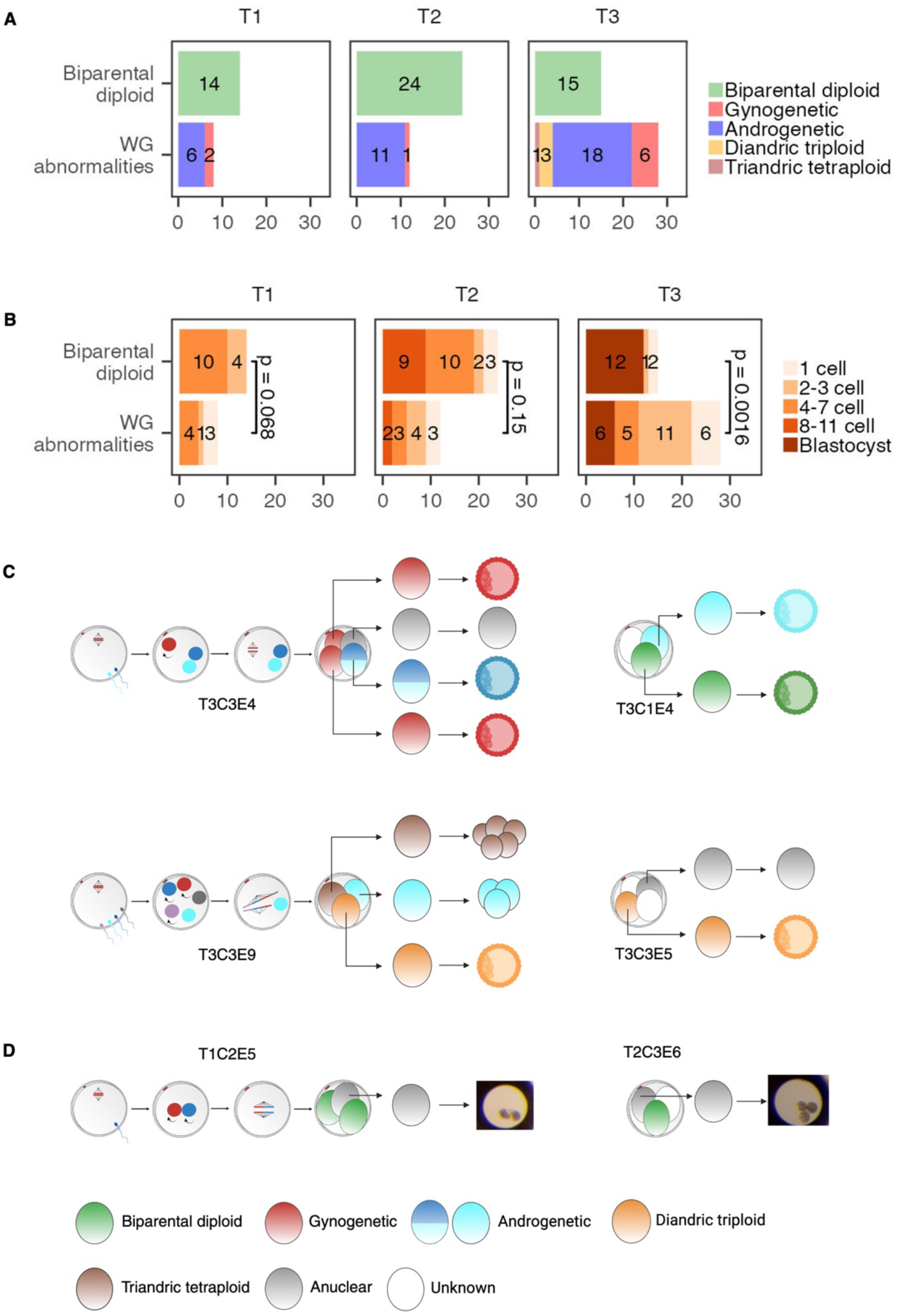
Blastomeres with whole-genome abnormalities can reach blastocyst stage despite impaired developmental potential. (A) Distribution of 53 biparental diploid blastomere outgrowths and 48 blastomere outgrowths with WG abnormalities across different timepoints of outgrowth collection. (B) Developmental stage comparison of biparental diploid blastomere outgrowths and blastomeres outgrowths with WG abnormalities in (A). (C) Schematic representation of the six blastomere outgrowths with WG abnormalities that reached the blastocyst stage at T3, along with their sibling blastomere outgrowths and original embryos. (D) Images of the two cleaved anuclear blastomere outgrowths and schematic diagrams illustrating the embryos from which they originated. In (C) and (D), the same symbols as in Figure 2C are employed. Fertilization and the first cleavage are not depicted for embryos containing blastomeres with an unknown haplotype (due to sample loss or haplotype failure).

Since aneuploidy is known to affect embryo development, we next evaluated the impact of heterogoneic division on aneuploidy levels in daughter blastomeres. By comparing the aneuploidy spectrum of blastomeres from both normal genome segregation and heterogoneic division, we observed a lower proportion of euploid blastomeres and a higher proportion of blastomeres with high-level aneuploidy (defined as >10% of autosome regions being aneuploid, with 10% corresponding to two large chromosomes) in blastomeres from heterogoneic division, compared to blastomeres originating from zygotes with normal genome segregation (**Supplemental Fig. S1**). Specifically, for blastomeres with WG abnormalities resulting from heterogoneic division, 32% (10 out of 31) were euploid, while 55% (17 out of 31) displayed high level aneuploidy. In contrast, for blastomeres from normal genome segregation, 56% (18 out of 32) were euploid, and only 3% (1 out of 32) exhibited high level aneuploidy (*p*=6.3×10^-6^) (**Supplemental Fig. S1**). Similarly, other studies demonstrated the increased rate of complex chromosomal abnormalities for embryos with multipolar division(Daughtry et al., 2019; Zhan et al., 2016). The association between heterogoneic division and vulnerability to chaotic chromosomal missegregations might be caused by the spindle abnormalities implicated in this process (Chatzimeletiou et al., 2005) **(Fig. 2C**). To control for the effect of aneuploidy on blastomere development, we compared the development of only euploid blastomeres. At T3, all biparental diploid blastomere outgrowths (100%,12 out of 12) reached blastocyst stage, while a much lower proportion (36%, 4 out of 11) of blastomere outgrowths with WG abnormalities reached blastocyst stage (*p*=0.0013). Additionally, for androgenetic blastomere outgrowths collected at T3, 2 out of 15 containing the X chromosome reached the blastocyst stage, while the 3 without the X chromosome did not, indicating that the Y chromosome alone is insufficient for blastocyst formation.

The majority of the 17 anuclear blastomeres arrested at the 1-cell stage. However, we identified 2 exceptions: one reached the 2-cell stage at T1, and the other reached the 3-cell stage at T2 (**Fig. 3D**), suggesting that the cytoplasmic divisions could occur without a nucleus. Additionally, we identified 16 more anuclear biopsies from 14 non-anuclear blastomere outgrowths at various stages of development, with the majority of the original blastomeres (12 out of 14) from embryos underwent multipolar first cleavage (**Supplemental Table S2**). Hence, the formation of anuclear blastomeres is not restricted to the first zygotic division. To confirm the origin of these empty outgrowths/biopsies, we examined the mitochondrial genotypes for empty biopsies obtained from the same cows. Maternal inheritance of the mitochondrial genome was corroborated by similar mitochondrial genotypes (**Supplemental Table S3**).

To conclude, blastomeres with WG abnormalities from heterogoneic division show diminished development. Despite this, certain uniparental/polyploid blastomeres still progress to the blastocyst stage (**Fig. 3C**), indicating their viability during the preimplantation stage and raising concerns about their further development and potential contribution to abnormal pregnancies.

### Whole-genome abnormalities do not alter the preimplantation developmental program but hinder transcriptomic development

We hypothesized that the impaired development of blastomere outgrowths with WG abnormalities would be reflected in the transcriptomic profiles of constituent cells, with WG abnormalities potentially causing deviations in the preimplantation developmental program. To investigate the effects of WG abnormalities on transcriptomic development, we conducted single-cell transcriptome sequencing on 677 cells derived from collected blastomere outgrowths. Among these, 446 transcriptomes met our quality control criteria, representing cells from 124 blastomere outgrowths derived from 49 embryos (**Supplemental Table S2; Methods**). We noted significantly higher fractions of mitochondrial transcripts in cells from blastocyst stage outgrowths compared to cells from other stages, except the 1-cell stage (**Supplemental Fig. S2**). The blastocyst stage cells with high fraction of mitochondrial transcripts were not enriched for cells with WG abnormalities. This finding suggests active mitochondrial gene expression in blastocysts and potential cytoplasmic RNA leakage in arrested 1-cell stage outgrowths. Consistent with our findings, previous studies utilizing PCR analysis have demonstrated a significant increase in mitochondrial transcripts at morula and blastocyst stages across various mammalian species(Ma et al., 2008; May-Panloup et al., 2005; Thundathil et al., 2005).

Following dimensionality reduction with Unified Manifold Approximation and Projection (UMAP), single-cell gene expression profiles exhibited a developmental trajectory from T1 to T3 (**Fig. 4A**). Cells with WG abnormalities did not form distinct clusters (**Fig. 4B**). Instead, by checking the expression of marker genes for essential preimplantation stages, we identified clusters corresponding to minor EGA, major EGA, inner cell mass (ICM), and trophectoderm (TE) stages, which we define as the molecular developmental stages of the cells. The cell cluster without EGA (pre-EGA) contained cells arrested before EGA (**Fig. 4C; Supplemental Fig. S3**). Pseudotime analysis with single-cell transcriptome data inferred developmental trajectories pointing to ICM and TE (**Fig. 4D**). These results indicate that transcriptomic changes are mainly driven by the natural developmental program rather than variations in genome constitution, aligning with previous single-cell transcriptome analyses of uniparental(Leng et al., 2019) and aneuploid(Fernandez Gallardo et al., 2023) cells from human preimplantation embryos.

**Figure 4.**
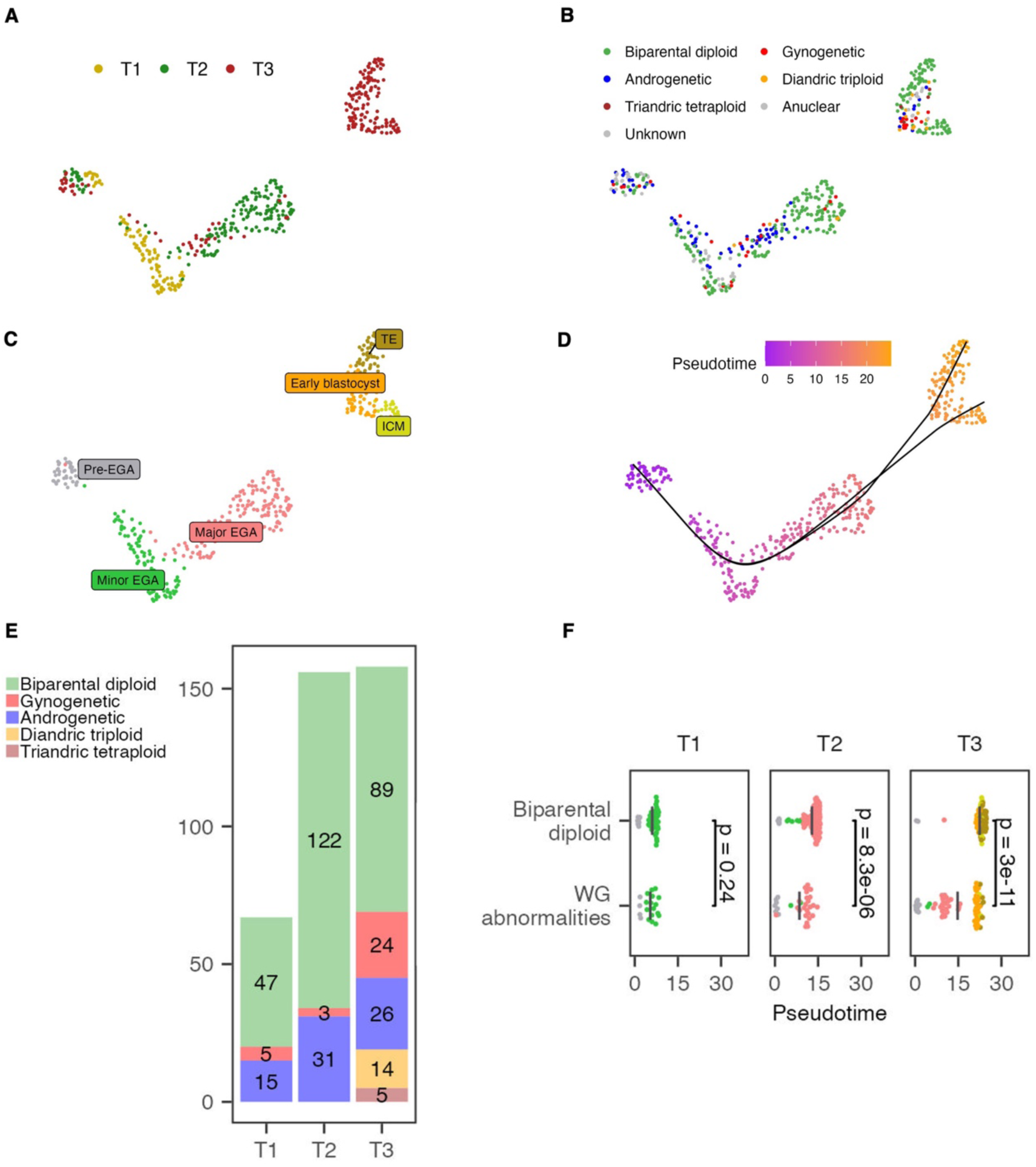
Whole-genome abnormalities do not alter the preimplantation developmental program but hinder transcriptomic development. UMAP for single-cell transcriptome data with cells colored by (A) time of outgrowth collection (B) genome constitution (C) molecular developmental stage indicated by marker genes expression (D) pseudotime value inferred from trajectory analysis. The inferred pseudotime trajectories are indicated by black curved lines on the UMAP. (E) Number of biparental diploid cells and cells with WG abnormalities used for transcriptome analysis for each timepoint. (F) Pseudotime comparison of biparental diploid cells and cells with WG abnormalities at each outgrowth collection timepoint. Each dot represents one cell, with the x-axis indicating its pseudotime value extracted from (D). Cells are colored according to their corresponding molecular developmental stage in (C) using the same color scheme. Vertical grey bars indicate the mean pseudotime value of each group.

We then explored whether the impaired development of blastomeres exhibiting WG abnormalities was reflected in single-cell transcriptome profiles. We characterized the physical age of the cells based on the time of outgrowth collection, while their molecular age was assessed using pseudotime values along the transcriptional trajectory. Compared to cells from biparental diploid outgrowths, those from outgrowths displaying WG abnormalities exhibited impaired development at T2 and T3, as evidenced by their smaller transcriptome-pseudotime ages (**Fig. 4E,F**; *p*=4.5×10^-5^ for T2 and *p*=4×10^-10^ for T3). This trend persisted when considering only cells inferred to be euploid (*p*=0.00026 for T2 and *p*=0.039 for T3). Our observation indicates the cell-level impaired transcriptomic development of blastomeres with WG abnormalities starting form T2, coinciding with major EGA.

Some blastomere outgrowths collected at T2 or T3 did not have the expected number of cells and appeared to be cleaving slower or blocked in their development. Surprisingly, when mapping the transcriptome profiles on the developmental trajectories, we observed that some cells from 1 cell and 2-3 cell stage outgrowths at T3 displayed profiles resembling major EGA, while some cells from 4-7 cell outgrowths exhibited profiles akin to TE or ICM (**Supplemental Fig. S4**). These observations suggest that transcriptomic development can continue in the absence of cell divisions.

### Distinct preimplantation cellular states are governed by specific key transcription factors

Since cells were clustered according to their molecular developmental stage regardless of genome composition, we next aimed to uncover the specific gene regulatory networks (GRNs) shaping cellular identity at each stage. We used pySCENIC to infer GRNs from single-cell RNA-seq data, based on co-expression of TFs and their target genes as well as cis-regulatory motif analysis. This approach identified 237 regulons, each comprising a transcription factor (TF) alongside its predicted target genes. Subsequently, we generated a UMAP visualization based on regulon activity, partitioning cells into distinct regulatory states (**Fig. 5A**). As anticipated, cells belonging to same molecular stage exhibited clustering, indicative of stage-specific regulatory dynamics. To further investigate the coactivity of TF combinations within each stage, we ordered cells based on their molecular stage and pseudotime values on regulon activity heatmap. The resulting plot revealed stage-specific regulatory states characterized by the activation or repression of specific TFs (**Fig. 5B**). We then selected key regulons with both high activity and specificity for each stage (**Fig. 5C; Supplemental Fig. S5; Methods**). Most of the TFs for selected key regulons have been shown to be crucial for embryonic development (**Supplemental Table S4**). For instance, GATA6 has demonstrated significance in the ICM of mouse, bovine, and human embryos(Marsico et al., 2023; Niakan and Eggan, 2013). Similarly, CDX2, GATA2 and GATA3 have been implicated in the TE of mouse, bovine and human embryos(Fernandez Gallardo et al., 2023; Gerri et al., 2020; Home et al., 2017; Nagatomo et al., 2013; Negrón-Pérez et al., 2017; Niakan and Eggan, 2013). Interestingly, for the pre-EGA stage, PRDM5 was identified as a key regulon, reflecting the arrested characteristics of these cells, as this TF is involved in G2/M arrest and apoptosis through suppressing the expression of several oncogenes and antagonizing WNT/β-catenin signaling(Deng and Huang, 2004; Shu et al., 2011). These findings shed light on the combinations of TFs that underlie cell state transitions and cellular identity during bovine preimplantation embryogenesis.

**Figure 5.**
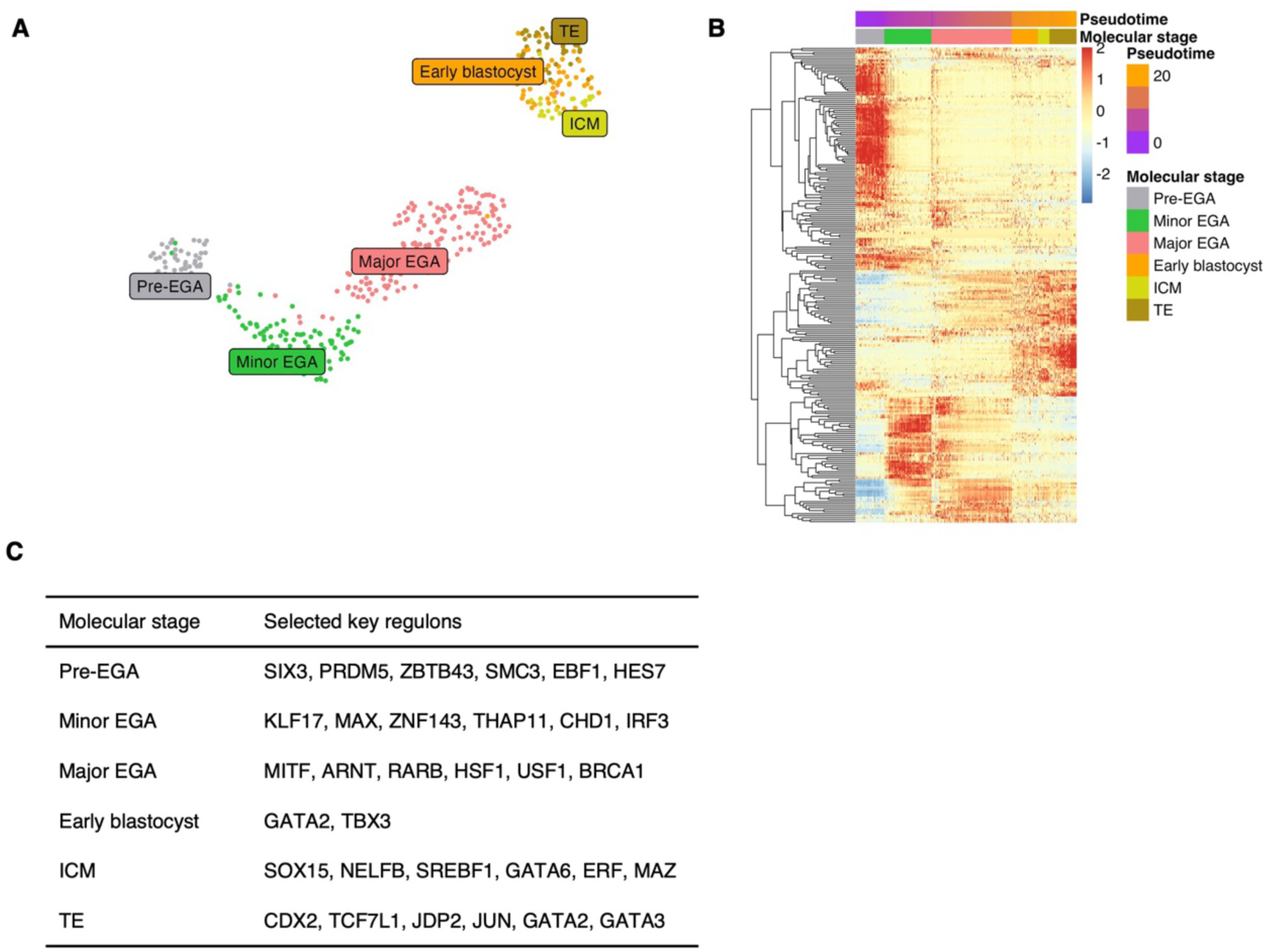
Distinct preimplantation cellular states are governed by specific key transcription factors. (A) UMAP displaying cell clustering based on regulon activity, with cells colored according to their macular developmental stage as depicted in Figure 4C. (B) Heat map of regulon activity. Each row represents one regulon and each column represent one cell. The plot is generated using rescaled Area Under the Curve (AUC) values, with cells ordered according to their molecular developmental stage and pseudotime values. (C) Key regulons selected for each stage.

### Whole-genome abnormalities induce stress responses during embryonic genome activation

Although WG abnormalities did not result in obvious deviations in the preimplantation developmental program, we hypothesized that the imbalanced parental genome compositions in cells with WG abnormalities could lead to parent-specific gene expression patterns for genes predominantly expressed from only one parent’s genome. Furthermore, these WG abnormalities may function as stress stimuli, inducing global transcriptomic changes that ultimately impair the overall developmental potential of cells. To investigate the parental genome-specific effects on gene expression, we compared the gene expression profiles of gynogenetic, androgenetic, and polyploid (diandric triploid and triandric tetraploid) cells to those of biparental diploid cells within each molecular developmental stage. Additionally, to capture the overall effects of WG abnormalities on gene expression, we aggregated all cells with WG abnormalities for combined analyses.

Differentially expressed (DE) genes were identified from the onset of EGA (**Fig. 6A**), suggesting that WG abnormalities lead to gene expression changes during EGA. Only one DE gene was identified at the early blastocyst stage, indicating a high degree of similarity between normal cells and cells with WG abnormalities just before the first lineage specification. DE genes found in the combined analysis partially overlap with those identified in separate analyses **(Supplemental Fig. S6; Supplemental Table S5)**. Manual functional annotation revealed some parental genome specific effects. We noted the downregulation of the maternally imprinted gene *SNRPN* in gynogenetic cells during major EGA, aligning with its known paternal-specific expression(Lucifero et al., 2006). Furthermore, we identified the up regulation of genes involved in cell adhesion (e.g., *RAPH1, PCDH17*) in polyploid cells during major EGA and at the TE stage (**Fig. 6B; Supplemental Table S5**). This indicates active cell adhesion-related processes, such as compaction in these cells, possibly caused by the additional paternal genome contribution.

**Figure 6.**
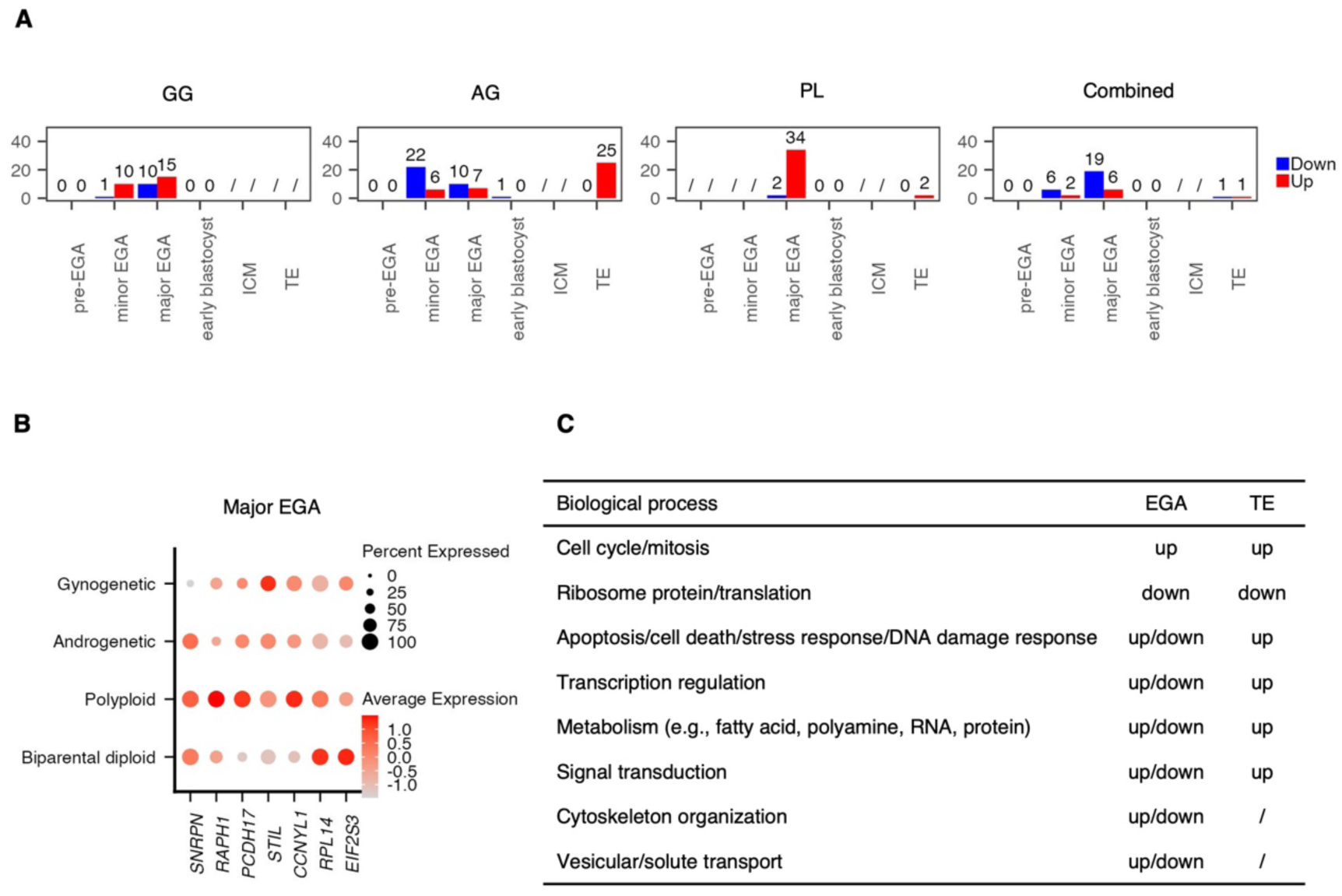
Whole-genome abnormalities induce stress responses during embryonic genome activation. (A) Numbers of DE genes identified in gynogenetic (GG), androgenetic (AG), polyploid (PL) and combined WG-aberrant cells compared to biparental diploid cells for each stage. The “/” symbol denotes comparisons not conducted due to insufficient cells. (B) Gene expression dot plot for example DE genes during major EGA. (C) Overall alterations in biological processes reflected by deviations in gene expression. “/” if no DE genes observed.

Other DE genes observed in uniparental/polyploid cells, as well as those identified through combined analysis, are mainly involved in similar biological processes, indicating global effects of WG abnormalities (**Fig. 6C; Supplemental Table S5**). We observed upregulation of cell cycle/mitosis-related genes (e.g., *STIL*, *CCNYL1*), indicating active preparation for cell cycle progression in cells with WG abnormalities. Meanwhile, we noted downregulation of ribosomal protein genes (e.g., *RPL14*) and translation initiation factor genes (e.g., *EIF2S3*). It is known that during cellular adaptation to stress stimuli, growth-related genes, including those involved in ribosomes and translation, are typically repressed, prioritizing resources towards stress protection over rapid proliferation(López-Maury et al., 2008). Our observation indicates reduced growth in cells with WG abnormalities as a consequence of the stress response triggered by WG abnormalities. Interestingly, for certain biological processes, DE genes were identified during EGA, leading to either upregulation or downregulation of these processes. No relevant DE genes were found for the early blastocyst stage, while in the TE, no DE genes or only DE genes resulting in upregulated activities of these processes were observed. The biological processes involved include apoptosis, cell death, stress response, DNA damage response, transcription regulation, metabolism, single transduction, cytoskeleton organization and transport **(Fig. 6C; Supplemental Table S5).** This indicates that cells with WG abnormalities adapt to stimuli caused by genome imbalance through transcriptional responses during EGA, involving transient upregulation or downregulation of relevant genes(López-Maury et al., 2008). A quiescent period follows during the early blastocyst stage, while the upregulation of specific genes in TE reflects establishment of new steady-state levels. Additionally, we conducted differential regulon analysis and DE regulons were identified for cells with WG abnormalities only during EGA (**Supplemental Table S5**). Most DE regulons are essential for embryonic development and their bidirectional changes mirror bidirectional gene expression alterations observed in differential gene expression analysis (**Supplemental Table S5**).

In summary, we observed parental genome-specific effects induced by WG abnormalities, suggesting distinct roles of the maternal and paternal genomes. Moreover, we identified universal cellular responses to WG abnormalities, marked by active preparation for cell cycle progression alongside reduced cell growth. Bidirectional alterations in specific biological processes and regulons during EGA, and their eventual balance, may dictate whether cells arrest or advance towards the blastocyst stage. Overall, these observed stress responses provide insights into the impaired development cause by WG abnormalities.

## Discussion

WG abnormalities are a type of chromosomal error frequently observed in spontaneous abortions(Hassold et al., 1980) and on very rare occasions in a chimeric state in live-born individuals(Madan, 2020). Contrary to its implications in abnormal pregnancy and congenital abnormalities, the proposed origins of such anomalies are deductions from surviving cells at later stages or remain speculative, and their effect on early development remains largely unknown. Here, we provide fundamental insights into the natural emergence and early developmental biology of gynogenetic, androgenetic, and polyploid blastomeres using haplotyping and single-cell transcriptome analysis. We demonstrated the spontaneous occurrence of uniparental and polyploid blastomeres through a non-canonical division during the first zygotic cleavage, which segregates parental genomes into distinct blastomeres. We not only confirmed our previous observation(De Coster et al., 2022) that heterogoneic division mostly coincides with multipolar first cleavage in polyspermic embryos, but also identified, for the first time, heterogoneic division occurring with bipolar first cleavage in normally fertilized embryos. The reduced developmental potential of uniparental and polyploid blastomeres derived from heterogonic division was evidenced by a lower blastocyst rate and a delay in pseudotime values beginning from T2, coinciding with major EGA. Stress responses observed during EGA suggests the cells’ efforts to combat fitness challenges induced by aberrant genome composition, with the outcome determining whether they arrest or progress to the blastocyst stage.

Although to our knowledge we are the first to explore the development of first-cleavage-derived uniparental and polyploid blastomeres, some studies have examined the preimplantation development of mammalian uniparental embryos. These embryos were generated through various methods, including nuclear manipulation(Hoppe and Illmensee, 1977; Leng et al., 2019; Xu et al., 2021), *in vitro* induction (Hao et al., 2004; Hwang et al., 2020; Ito et al., 1991; Leng et al., 2019; Liu et al., 2002) or natural occurance(Hardy et al., 1989) of parthenogenetic development in unfertilized oocytes, as well as bisection of one-cell fertilized eggs (Tarkowski, 1977). Aligned with the impaired developmental potential observed in blastomeres with WG abnormalities in our study, those studies on preimplantation uniparental embryos have shown a decrease in the morula/blastocyst rate compared to biparental diploid embryos., alongside poorer morphology and reduced cell count in resulting blastocysts(Hardy et al., 1989; Leng et al., 2019). Several studies have monitored post-implantation development and affirmed the compatibility of diploid uniparental genome constitution with later development. For instance, porcine parthenogenetic fetuses can develop until day 35 with abnormal organ morphologies (Hwang et al., 2020). Additionally, live-born fertile mice have been created via both androgenesis and gynogenesis(Hoppe and Illmensee, 1977). Moreover, diploid gynogenetic cells have been identified in various organs of live-born chimeric mice developed from aggregated chimeric embryos(Ito et al., 1991). These findings are consistent with observations in humans, suggesting that uniparental cell lines are typically selected against during development but can persist until late pregnancy and even be compatible with live birth, raising various concerns. Our study demonstrates that heterogoneic division could serve as a spontaneous origin of uniparental and polyploid cell lines observed during late development.

The parental genome specific effects identified in this study correlates with findings from a prior investigation of uniparental human embryos(Leng et al., 2019). As for the stress response observed solely in our study, there are possible causes aside from species-specific effects and sample variations(Xu et al., 2021). The referenced study created diploid gynogenetic and androgenetic embryos through nuclear manipulation and induction of parthenogenesis, respectively. Only embryos demonstrating good morphology and developmental speed were utilized. Only XX and XY androgenetic embryos were included. And their analysis focused exclusively on genes with highly biased expression in gynogenetic or androgenetic embryos. In contrast, our study focused on a broad spectrum of spontaneously occurring WG abnormalities, which frequently exhibit complex aneuploidy and contain either haploid or diploid genome constitutions. While haploidy has been demonstrated to convert frequently to diploidy during development(Ito et al., 1991; Leng et al., 2017; Tarkowski, 1977), both complex aneuploidy and haploidy are known to impact preimplantation development(Liu et al., 2002; McCoy et al., 2015). Additionally, our cohort included androgenetic blastomeres lacking the X chromosome, a condition known to cause embryonic arrest. Finally, rather than focusing solely on genes with highly biased expression in gynogenetic or androgenetic embryos, our study examined the broader impact of WG abnormalities on global gene expression changes. These factors led us to observe EGA as a critical period for embryos with WG abnormalities. During this stage, their constituent cells exhibit stress responses in gene expression, which could determine whether they arrest or successfully progress to the blastocyst stage.

We identified anuclear cells from outgrowth at different stages, which is in agreement with previous observations in rhesus(Daughtry et al., 2019) and human embryos(Hardy et al., 1993). Consistent with our previous analysis(De Coster et al., 2022), most of the anuclear blastomeres were identified from blastomere outgrowths following multipolar first cleavage, suggesting a potential association between the first cleavage pattern and the origin of these anuclear cells. In addition, we demonstrate that some of the anuclear cells can cleave without nuclei. Cytoplasmic divisions are generally thought to rely on nuclear divisions and mitotic signals. However, a recent study by Bakshi et al. observed that cytoplasm can compartmentalize, and a rare fraction of the compartments can divide repeatedly without nuclei and independent of mitotic CDK/cyclin complexes during normal drosophila embryogenesis (Bakshi et al., 2023). They showed that this phenomenon is relevant to mitotically delayed nuclei. We hypothesize that a similar mechanism may apply here, with the anuclear cells derived from mother cells containing nuclei with delayed mitosis. These findings collectively underscore the need for further investigation into the detailed mechanisms underlying the presence of anuclear cells in mammalian embryos and their potential roles in embryo development.

Insights from this study raise clinical concerns. Firstly, in IVF laboratories, human embryos with two pronuclei (PN) are preferentially selected for transfer(Balaban et al., 2011). We demonstrate here that zygotes with three PN may result in a mixture of biparental diploid and uniparental/polyploid blastomeres. Since the latter are developmentally compromised, the resulting embryo may still develop normally. Secondly, there is a growing trend in many laboratories to monitor embryonic development using time-lapse microscopy and rank embryos based on cleavage kinetics(Balaban et al., 2011; Sigalos et al., 2016; Sugimura et al., 2017, 2012). Embryos with multipolar first cleavage are typically given a low rank. We showed here that 22% of embryos with a multipolar first cleavage exhibited normal, equal division of the genome with additional anuclear cells. It would be unjust to low rank these embryos since they have the potential to lead to healthy babies. Additionally, even though uniparental/polyploid cells are normally generated with multipolar first cleavage, it is not uncommon that biparental diploid blastomeres are generated together, which may gain developmental advantage and lead to normal development. Thirdly, embryos typically ranked high, characterized by a normal pronuclei count and bipolar first cleavage, may still experience WG segregation errors, leading to the generation of solely uniparental cells, posing a potential risk of abnormal pregnancy. Rather than removing or keeping embryos based on apparent abnormalities, it might be better to evaluate the genome constitution of preimplantation embryos and deselect those with WG abnormalities, while retaining those with a normal chromosome count and parental haplotype constitution. Preimplantation genetic testing for aneuploidy (PGT-A) has become a common practice to select embryos with a normal number of chromosomes for transfer. However, the majority of PGT-A tests only measure aneuploidies, with only some measuring the parental haplotype constitution(Coonen et al., 2020). We suggest the use of genotyping/haplotyping together with PGT-A(Masset et al., 2022; Zamani Esteki et al., 2015) to check both chromosome count and parental haplotype constitution and select against WG abnormalities. This approach is likely to further improve the overall IVF success rate(Caroselli et al., 2023).

In conclusion, this study provides comprehensive insights into the natural origin and early development of uniparental/polyploid blastomeres. We demonstrated their overall impaired developmental potential, primarily due to stress responses triggered by WG abnormalities during EGA. Blastomeres that successfully navigate through this phase and progress to the blastocyst stage may constitute a significant fraction of embryos and contribute to abnormal development. Our findings underscore the importance of routine screening against embryos bearing WG abnormalities in IVF labs.

## Materials and Methods

### Single-cell collection from bovine blastomere outgrowths

#### In vitro embryo production

Prior ethical agreement was not necessary. Ovaries from cows raging 4 to 5 years were collected post-mortem in a commercial slaughterhouse. The sperm used for *in vitro* fertilization was collected from a 5-years-bull. Standard in vitro procedures were followed to produce bovine embryos (Wydooghe et al., 2014). Briefly, bovine (*Bos taurus*) ovaries were collected and processed within 2 h. The ovaries were rinsed three times in warm physiological saline supplemented with 0.25µg/mL kanamycin. Using an 18-G needle and a 10-mL syringe, antral follicles (2 – 8 mm diameter) were punctured and kept separately per ovary in 2.5 mL Hepes-Tyrode’s albumin-pyruvate-lactate (Hepes-TALP). Using a stereomicroscope, cumulus-oocyte complexes were collected and washed in Hepes-TALP and then in maturation medium (modified bicarbonate-buffered TCM-199 supplemented with 50 ppm gentamicin and 20 ng/mL epidermal growth factor). Cumulus-oocyte complexes were *in vitro* matured per donor in four-well dishes (NuncTM) in 500 μL maturation medium for 22 h at 38.5°C in 5% CO_2_ in humidified air. *In vitro* fertilization was performed with frozen-thawed semen from a Holstein-Friesian bull (*Bos taurus*) after selection over a discontinuous 45/90% Percoll^®^ gradient (GE Healthcare Biosciences, Uppsala, Sweden). The mature oocytes were fertilized by incubating them with spermatozoa at a concentration of 1 × 10^6^ spermatozoa/mL in IVF–TALP medium enriched with BSA (Sigma A8806; 6 mg/mL) and heparin (20 μg/mL) for 21 hours at 38.5°C, 5% CO_2_ in humidified air. After fertilization, the presumed zygotes were vortexed in 2.5 mL HEPES-TALP for 3 minutes to remove the cumulus and sperm cells adhered to the *zona pellucida* and subsequently transferred to 50 μL droplets of synthetic oviductal fluid (SOF), enriched with 4 mg/mL BSA (Sigma A9647), non-essential and essential amino acids (SOFaa), 5 μg/mL insulin, 5 μg/mL transferrin, and 5 ng/mL selenium. The droplets were covered with 900 μL paraffin oil (SAGE oil for tissue culture, ART-4008-5P, Cooper Surgical Company). *In vitro* culture was performed at 38.5 °C in 5% CO_2_, 5% O_2_, and 90% N_2_.

#### Blastomere dissociation and culture

The presumptive zygotes were monitored from 26 to 36 hours post-fertilization (hpf) every 30 minutes to identify a direct cleavage of the zygote into three or four blastomeres (multipolar division) or into two blastomeres (bipolar division). Blastomeres from embryos that cleaved into more than four cells or presented multiple fragments were not processed further. Immediately upon the first division, embryos were washed in Hepes-TALP and treated with 0.1% pronase (protease from *S. griseus*) in TCM-199 to dissolve the *zona pellucida*. Next, the embryos were washed in TCM-199 with 10% FBS and then transferred to Ca^+2^/Mg^+2^-free PBS with 0.05% BSA to enhance blastomere dissociation, which was performed using a STRIPPER pipet holder and 170 μm and 135 μm capillaries (Origio, Cooper Surgical, CT, US) in Ca^+2^/Mg^+2^-free PBS supplemented with 0.1% polyvinylpyrrolidone (PVP). Subsequently, single blastomeres were washed in culture medium and transferred individually to one well of a Primo VisionTM micro well group culture dish (Vitrolife, Göteborg, Sweden), which contained a total of 16 small wells covered by a 40 μL droplet of culture medium and 3.5 mL paraffin oil. Single blastomeres were cultured at 38.5°C in 5% CO_2_, 5% O_2_, and 90% N_2_ until 72, 121 or 170 hpf according to the experimental design.

#### Single-cell isolation

For isolation of single cells at 72 hpf, 121 hpf or from blastomeres that did not reach the blastocyst stage at 170 hpf, the outgrowths were transferred to Ca^+2^/Mg^+2^-free PBS with 0.1% PVP, and the cells were dissociated mechanically with a STRIPPER pipet holder and 170 μm and 135 μm capillaries (Origio, Cooper Surgical, CT, US). For blastomeres that reached the blastocyst stage at 170 hpf, single-cell isolation was done by incubating the embryos in trypsine-EDTA at 38.5°C, followed by washing in Ca^+2^/Mg^+2^-free PBS with 0.1% PVP, pipetting with a STRIPPER pipet holder, and 135 μm and 70 μm capillaries (Origio, Cooper Surgical, CT, US) and micromanipulation with holding (MPH-MED-35, Origio, Cooper Surgical, CT, US) and biopsy (MBB-BP-M-30, Origio, Cooper Surgical, CT, US) pipettes. When single-cell dissociation was not possible, clusters of cells were collected. Single cells from all time points were washed in Ca^+2^/Mg^+2^-free PBS with 0.1% PVP before transferring them into a well of a skirted 96-well plate (4ti-0960/C, AZENTA Life Sciences, Bioké, Leiden, The Netherlands) containing 2.5 µL of RLT lysis buffer with a 70 µm capillary (Origio, Cooper Surgical, CT, US) and a STRIPPER pipet holder screwed on 0.5 µL. The collected samples were kept on ice during the whole procedure and then stored at −80°C.

### DNA and RNA separation, library preparation and sequencing

#### Separation of DNA and RNA from single cells

The DNA and mRNA from single cells were separated following the genome and transcriptome sequencing (G&T-seq) protocol(Macaulay et al., 2016) on a robotic liquid-handling platform (Microlab STAR Plus, Hamilton). Specifically, the 96-well sample plate was positioned on the robot deck alongside (a) a plate for capturing poly-A mRNAs, containing 10µl per well of Dynabeads^®^ MyOneTM Streptavidin C1 (Thermo Fisher Scientific) bound to biotinylated poly-dT oligos containing the SmartSeq2 primer sequence ‘5BioTinTEG/ - AAGCAGTGGTATCAACGCAGAGTACTTTTTTTTTTTTTTTTTTTTTTTTTTTTTTVN’ (IDT), (b) a plate for washing away DNA from the cell lysate supernatant, containing 25µl per well of G&T-wash buffer, and (c) an empty DNA destination plate. First, the cell lysate was mixed with biotinylated poly-dT beads and incubated for 20 min. Subsequently, beads bound to mRNA were pulled down using a low elution magnet (Alpaqua) for 2 min, and the supernatant was transferred to the DNA destination plate. Following this, the beads were subjected to two washes, each with 10µl of G&T wash buffer, and the supernatant was transferred to the DNA destination plate. The DNA destination plate, containing 37.5µl of G&T wash buffer, was then centrifuged for 1 min at 1000g and stored at -80°C.

#### RNA amplification, library preparation and sequencing

The plate containing poly-A mRNA-bound beads was processed using an adapted SmartSeq2 protocol with 20 PCR cycles. Subsequently, the amplified single-cell cDNA was purified using a 0.8:1 ratio of Agencourt AMPure XP beads (Analis), followed by washing with 80% ethanol and elution in water. Libraries were generated from the amplified cDNA according to the Nextera XT (Illumina) protocol with quarter volumes. These libraries were pooled and sequenced using single-end 50 sequencing on a HiSeq4000 Illumina sequencer, aiming at 1 million reads per sample. For a subset of blastocyst stage single cells, the libraries were resequenced paired-end 150 on a NovaSeq 6000 Illumina sequencer for more reads.

#### DNA amplification, GBS library preparation and sequencing

For each blastomere outgrowth, DNA from either individual single cell or several cells was used for GBS (with double enzyme restriction) processing. A sibling blastocyst was collected and processed together when available. Whole-genome amplification (WGA) was carried out using the REPLI-g SC kit (Qiagen, Germany) following the manufacturer’s guidelines, with variations in reaction volumes (full or half) and an incubation period of 2-3 hours. The DNA separated from the single cells stored in G&T wash buffer (see before), was thawn on ice, centrifuged for 1 min at 1000g, purified with Agencourt AMPure XP beads (Beckman Coulter, USA), eluted in 4 µl of scPBS and processed following the REPLI-g SC kit (Qiagen, Germany) manufacturer’s guidelines with an incubation time of 2h. Additionally, bulk DNA was extracted from ovarian tissue of the donor cows (mothers of the respective embryos), semen from the bull (father of the embryos), and blood from the parents of the bull (paternal grandparents of the embryos) using the DNeasy Blood and Tissue kit (Qiagen, Germany).

The whole-genome amplified DNA and bulk DNA were subjected to double restriction digestion with 8 units (U) of PstI-HF (R3140S, NEB) and 4U of CviAII (R0640L, NEB) enzymes (New England Biolabs, NEB, USA) combined with adapter ligation using 200-300 ng as an input DNA for each sample. The restriction-ligation reaction was performed in a total reaction volume of 15 µl, with a final concentration of 1X rCutSmart buffer (B6004S, NEB), 1mM ATP (P0756S, NEB) and 160U of T4 DNA Ligase (M0202L, NEB). Next, the double-sided size-selection was performed with Agencourt AMPure XP beads (Beckman Coulter, USA) and 7 cycles of PCR to amplify and barcode size-selected adapter-ligated fragments using Q5 High-Fidelity 2X Master Mix (NEB) and 0.5 µM primer mix with the following program. Subsequently, the libraries were purified with Agencourt AMPure XP beads (Beckman Coulter, CA, USA), equimolarly pooled and sequenced paired-end 150 on NovaSeq 6000 Illumina sequencer, with a target of 20-30 million reads per sample.

### Data analysis

#### GBS data processing, haplotyping and aneuploidy profiling

The raw reads were processed using fastp (v0.23.2)(Chen et al., 2018) to trim adaptor sequences, filter out low-quality reads, remove UMI sequences and append them to read names. Next, cutadapt (v 1.18)(Martin, 2011) was applied to select reads that start with the internal barcode and trim off the barcode. Reads were then mapped to bovine reference genome bosTau9 (ARS-UCD1.2) using BWA-MEM (v0.7.17)(Li, 2013). Next, UMICollapse(Liu, 2019) was used for collapsing duplicated reads with the same UMI, while accounting for sequencing/PCR errors. Across all samples, we obtained a median of 64 million mapped reads. After UMI deduplication, a median of 35 million mapped reads were retained (**Supplemental Table S2**). Variant calling was performed with freebayes (v 1.3.2)(Garrison and Marth, 2012). Initially, variants were called for the parents and phasing reference(s) (parental grandparents or a sibling blastocyst). Subsequently, the identified variants were utilized as input to call variants for each single/multi-cell sample. The called SNVs were converted into bi-allelic calls (e.g., AA, AB, and BB) with B-allele frequency (BAF) values calculated based on allele-specific depth of coverage using a custom R script. The genotype calls and BAFs were then used as input for the siCHILD pipeline (details in the method paper(Zamani Esteki et al., 2015)). In brief, siCHILD performs pedigree-based haplotyping analysis. The parents were first phased using phasing reference(s). Subsequently, for specific combinations of phased parental genotypes, corresponding SNP BAF values of the sample of interest were retrieved and plotted on paternal and maternal haplarithms. Genome-wide parental haplotype inheritance can then be inferred through visual inspection of the genome-wide haplarithm plots.

In addition to haplotype information, the R package QDNAseq and custom scripts were applied to BAM files post-deduplication for aneuploidy profiling with a fixed bin size of 100kb. To quantify the extent of aneuploidy, the percentage of aneuploid autosomal regions was calculated for each sample. Specifically, bins with a segmentation logR value within *log*2(1/2)) and log2(3/2) were classified as normal, while all other bins were classified as abnormal. The percentage of aneuploid autosomal regions was calculated by summing up the length of all abnormal bins and dividing it by the size of all autosomes. Samples were categorized based on the percentage of aneuploid autosomal regions: ≤ %1 as euploid, 1-5% as low-level aneuploidy, 5-10% as medium-level aneuploidy, and >10% as high-level aneuploidy. To determine chromosome X nullisomy status, the lengths of all bins on chromosome X with a segmentation logR value ≤ −3 were aggregated. The fraction of the chromosome X region exhibiting nullisomy was determined by dividing this sum by the size of chromosome X. Samples with >95% nullisomy in the chromosome X region were classified as chromosome X nullisomy.

The haplotype composition and aneuploidy status of each blastomere outgrowth were determined using GBS results from a single-cell/multi-cell sample. A sample was considered anuclear if >60% and >10% of the reads mapped to the mitochondrial genome before and after UMI deduplication, respectively, with empty haplarithm and copy number plots. To infer an outgrowth to be anuclear, each cell in the outgrowth was individually checked and confirmed to be anuclear. To verify the presence of mitochondrial genomes in inferred anuclear biopsies and ascertain their maternal inheritance, we exclusively called variants on the mitochondrial genome across all inferred anuclear biopsies using Freebayes (v1.3.2)(Garrison and Marth, 2012) and compared the called genotypes (**Supplemental Table S3**).

#### Single-cell RNA sequencing data preprocessing

The raw reads were processed with fastp (v0.23.2)(Chen et al., 2018) to trim adaptor sequences and filter out low-quality reads. Subsequently, the reads were mapped to the bovine reference genome bosTau9 (ARS-UCD1.2) using STAR aligner (v 2.7.3)(Dobin et al., 2013) with Ensembl annotations ARS-UCD1.2.105. Next, raw counts per cell were obtained using HTSeq (v 0.9.1)(Anders et al., 2015). Cells with fewer than 200,000 detected molecules or fewer than 2,000 expressed genes were excluded from further analysis, resulting in 446 cells out of 677 passing these quality control thresholds. The retained cells exhibited a median of 862,384 molecules and 7,272 genes detected per cell (**Supplemental Table S2**). Counts were then normalized and scaled using R package Seurat (v5.0.1)(Hao et al., 2023).

#### Cell clustering, cell type identification and trajectory analysis

Cell clustering was performed based on the normalized and scaled transcriptome profiles using UMAP implemented in Seurat (v5.0.1). Subsequently, we assigned molecular cell stage labels to each cluster based on the expression of (a) known marker genes for ICM and TF (e.g. FN1, KDM2B for ICM, GATA3 for trophectoderm, etc.(Nagatomo et al., 2013; Negrón-Pérez et al., 2017)); (b) genes first expressed at 4-cell stage (minor EGA) or 8-16 cell stage (major EGA) in bovine embryos (Graf et al., 2014). For trajectory analysis, we employed the R package Slingshot (v2.10.0) to infer developmental trajectories based on transcriptome profiles.

#### Regulon analysis

Regulon analysis was performed on the transcriptome data using pySCENIC (v0.12.1)(Van de Sande et al., 2020). First, for each motif in the public collection (https://resources.aertslab.org/cistarget/motif_collections/), genes in the bovine reference genome bosTau9 (ARS-UCD1.2) with orthologs in humans were prioritized. This prioritization was determined by a scoring system that assessed the presence of the motif within a 10kb window upstream and downstream of the transcription start site. Consequently, a motif x gene matrix was generated. Next, this matrix served as input alongside the raw count matrix to pySCENIC (v0.12.1) to detect active regulons. pySCENIC was run ten separate times, and only regulons appearing two or more times were retained. The final AUC matrix was constructed using the maximum AUC values for each regulon among all runs in which they were present. The AUC heatmap was created with AUC values scaled for each regulon. To rank the identified regulons according to their specificity in each cell lineage, a regulon specificity score was calculated using “calcRSS” function within R package SCENIC (v 1.3.1). To identify the key regulons for each molecular stage, we considered only regulons where corresponding TFs were expressed in more than 50% of the cells. From these, for each stage, we selected the top 6 regulons with the most frequent TF expression among those ranked in the top 20 for both activity and specificity.

#### Differential expression and regulon analysis

Differential expression analysis was performed by comparing androgenetic/gynogenetic/polyploid cells to biparental diploid cells within each molecular cell stage using the FindMarkers function in Seurat (v5.0.1), employing the “MAST” method with a minimum percentage threshold of 0.5. Pseudotime values and aneuploid status were incorporated as latent variables in the model. Genes were considered differentially expressed if they met the criteria of an adjusted *p*-value < 0.05 and an absolute log2 fold change greater than 1. Differential regulon analysis was performed between cells exhibiting WG abnormalities and biparental diploid cells within each molecular cell stage by Wilcoxon ranked-sum test on the AUC values. Regulons were considered to have significantly different activity if they demonstrated an adjusted *p*-value < 0.05 and an absolute log2 fold change greater than 1. The functions of the differentially expressed genes and regulons were curated through manual online searches and literature reviews.

#### Statistical analysis

Categorical data were compared using the chi-squared test, with Fisher’s exact test applied when cell counts were too small. Pseudotime values were compared using the t-test. Statistical significance was determined using a two-tailed approach with a significance threshold of *p* < 0.05. All tests were conducted in R (v4.3.2).

## Data availability

The GBS and single-cell RNA sequencing data reported in this paper are available at the European Nucleotide Archive (ENA) under project number PRJEB76932 (https://www.ebi.ac.uk/ena/browser/view/PRJEB76932).

## Acknowledgements

Funding was received from the Marie Skłodowska-Curie grant agreement No 813707 (MATER) and from the KU Leuven, C1-C14/22/125 to J.R.V. and T.V. T.L. and T.V. were supported by the Research Foundation Flanders (FWO: G0C6120N, G088621N and I001818N). Y.Z. was supported by the Marie Skłodowska-Curie grant agreement No 813707 (MATER). A.F. is supported by the European Union’s Horizon 2020 research and innovation programme under the Marie Skłodowska-Curie grant agreement No 860960 and BOF22/ITN/036.

