## Supplemental Figure S1-S6 for "Origin and development of uniparental and polyploid blastomeres"

### Slide 1
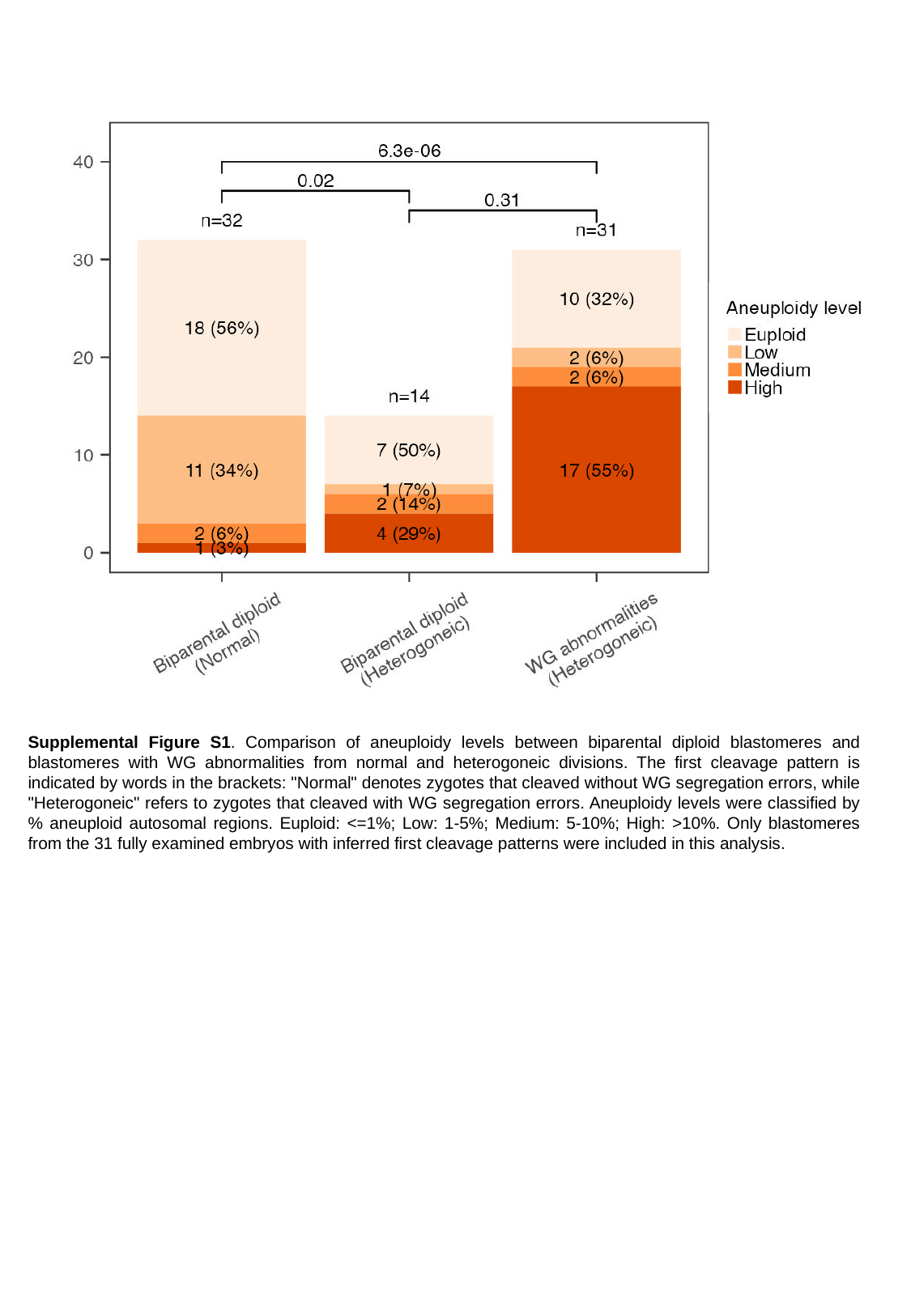

Supplemental Figure S1. Comparison of aneuploidy levels between biparental diploid blastomeres and blastomeres with WG abnormalities from normal and heterogoneic divisions. The first cleavage pattern is indicated by words in the brackets: "Normal" denotes zygotes that cleaved without WG segregation errors, while "Heterogoneic" refers to zygotes that cleaved with WG segregation errors. Aneuploidy levels were classified by % aneuploid autosomal regions. Euploid: <=1%; Low: 1-5%; Medium: 5-10%; High: >10%. Only blastomeres from the 31 fully examined embryos with inferred first cleavage patterns were included in this analysis.

### Slide 2
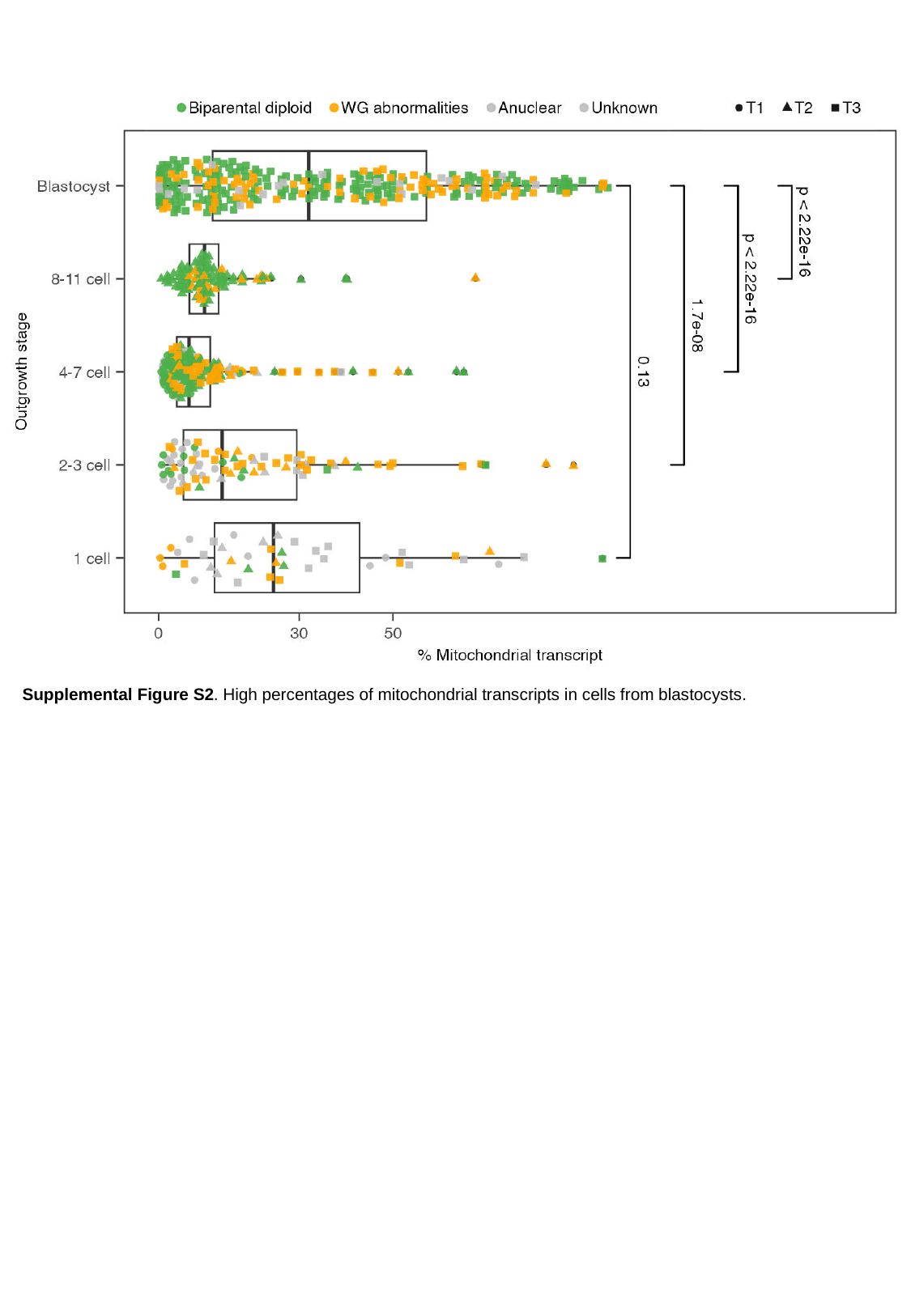

Supplemental Figure S2. High percentages of mitochondrial transcripts in cells from blastocysts.

### Slide 3
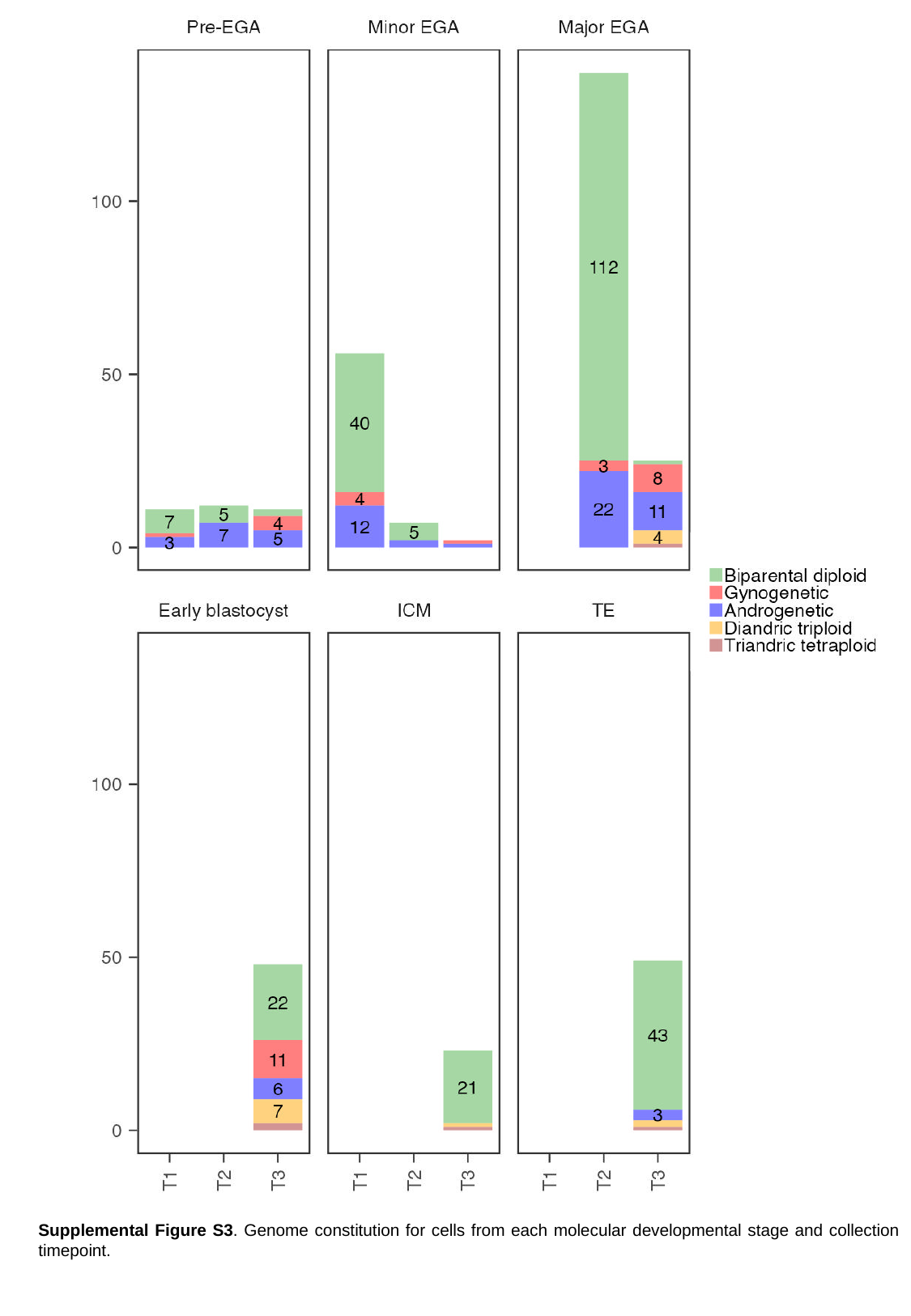

Supplemental Figure S3. Genome constitution for cells from each molecular developmental stage and collection timepoint.

### Slide 4
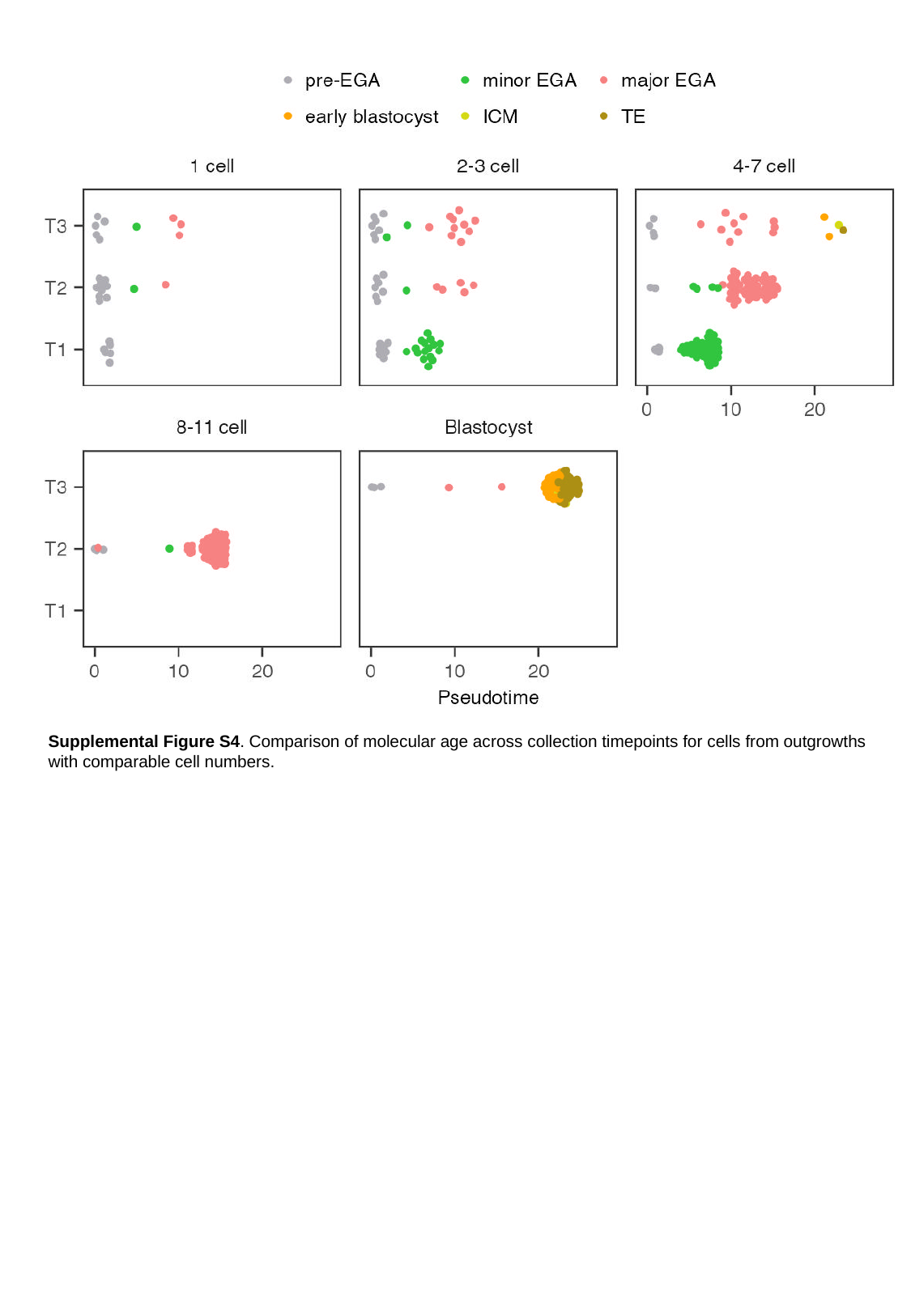

Supplemental Figure S4. Comparison of molecular age across collection timepoints for cells from outgrowths with comparable cell numbers.

### Slide 5
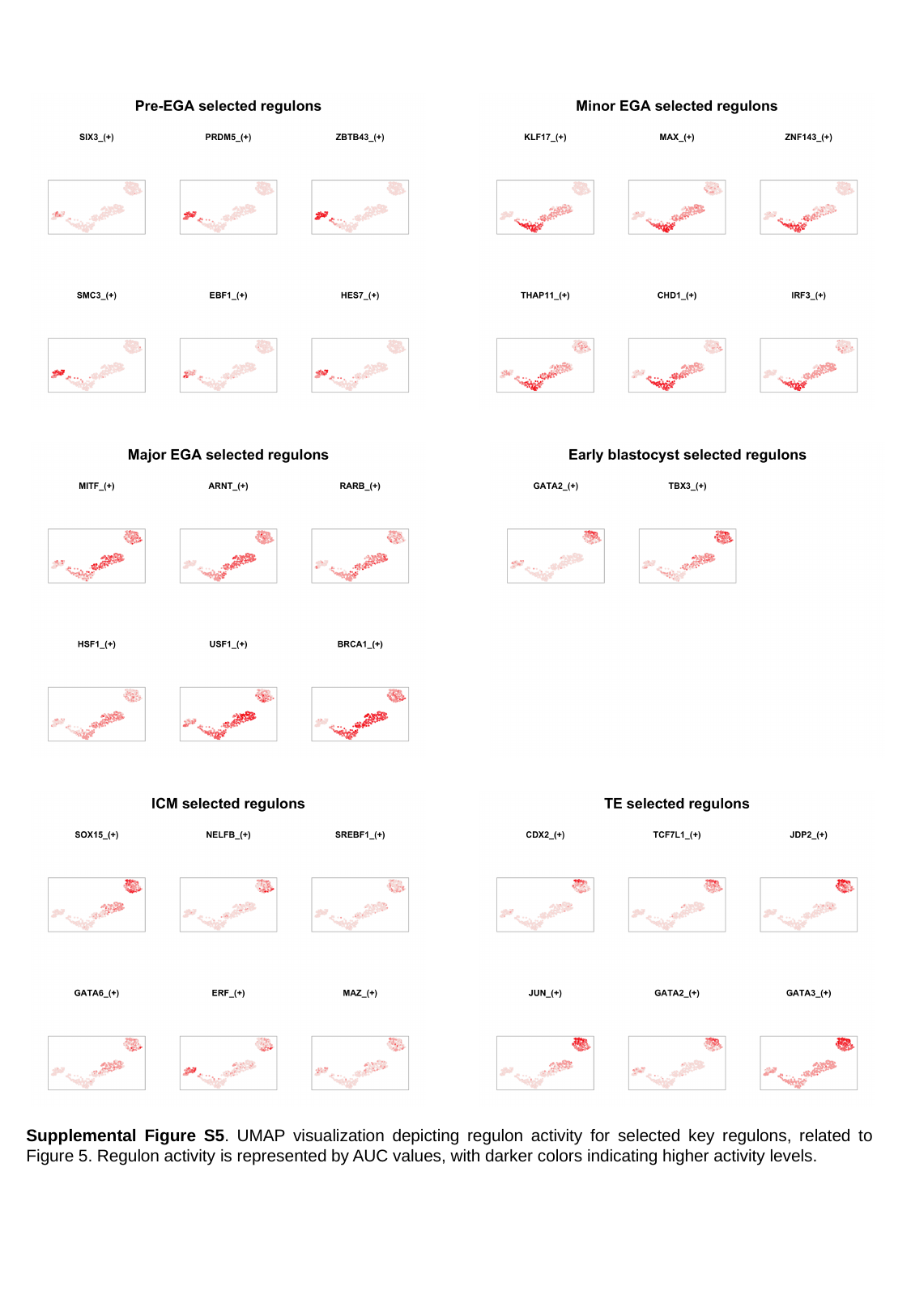

Supplemental Figure S5. UMAP visualization depicting regulon activity for selected key regulons, related to Figure 5. Regulon activity is represented by AUC values, with darker colors indicating higher activity levels.

### Slide 6
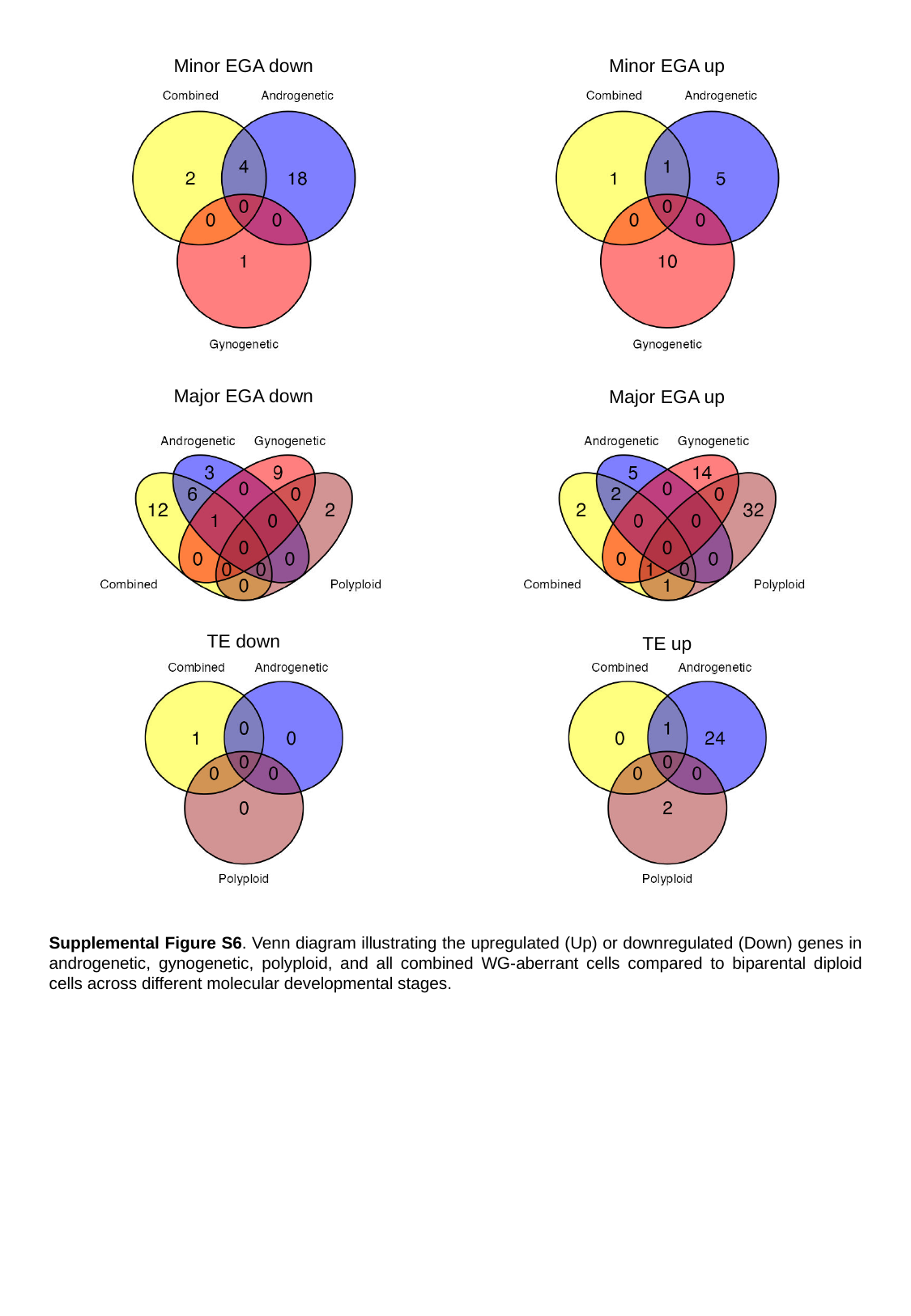

Minor EGA down
Minor EGA up
Major EGA down
Major EGA up
TE down
TE up
Supplemental Figure S6. Venn diagram illustrating the upregulated (Up) or downregulated (Down) genes in androgenetic, gynogenetic, polyploid, and all combined WG-aberrant cells compared to biparental diploid cells across different molecular developmental stages.
